# SARS-CoV-2 spike protein-associated sialoglycoconjugates induce nanoscale filipodia to facilitate micro-size platelet clotting

**DOI:** 10.64898/2026.04.19.719510

**Authors:** Ablikim Baqi, Ainur Sanaubarova, Belinda L Spillings, Erwan Bremaud, Veronika Masic, Larissa Dirr, Mark von Itzstein, Johnson Mak

## Abstract

COVID-19 disease is associated with thrombosis, but the pathogenic mechanism remains unclear. Here, we investigate how SARS-CoV-2 spike protein causes platelet activation and aggregation. Our three-dimensional ultrastructural analyses showed that invaginated platelet structures, open canalicular system (OCS), expanded upon activation, trapping viral particles in the process. Binding with platelet OCS concealed SAR-CoV-2 spike-coated particles from virion detection in platelet-depleted blood plasma. Both SARS-CoV-2 spike coated-particles and recombinant spikes specifically induced platelet aggregation with nanoscale filipodia extensions, with the terminal sialic acids of the SARS-CoV-2 spike protein-associated sialoglycoconjugates being the key determinant in platelet activation. Our work illustrates that virus-associated sialic acids, not proteins, are functionally responsible for SARS-CoV-2 induced thrombotic events, providing a mechanistic insight on how glycosylation contributes to disease severity in COVID-19. This study lays the foundation for the development of glycan-modified vaccines with reduced risks of thrombosis.

## Introduction

Coagulopathy-associated microvascular thrombosis and vascular dysfunction are hallmarks of hospitalised patients with severe acute respiratory syndrome coronavirus (SARS-CoV-2) disease ^1–3^, in which high incidence of thromboembolic events is found in autopsy of deceased coronavirus disease 2019 (COVID-19) patients ^4^. As SARS-CoV-2 is more difficult to detect in blood than in nasopharyngeal swabs ^5^, platelet-associated haemostatic abnormalities in COVID-19 are thought to be a secondary consequence of inflammation or cytokine storm from SAR-CoV-2 infection ^6–9^. SARS-CoV-2 effectively infect endothelial cells ^1^, while platelets amplify the endotheliopathy of SARS-CoV-2 thereafter ^10^. Despite the benefit of SARS-CoV-2 vaccination outweighing the diseases-associated risks ^11–13^, vaccine-related thrombotic events occur rarely at 0.21 cases per 1 million COVID-19 vaccinated person-days ^14^. Amongst these thrombotic events, a sub-group of this thrombotic disorder is known as vaccine-induced immune thrombotic thrombocytopenia (VITT) that is associated with pathogenic anti-platelet factor-4 (anti-PF4) antibodies ^15–17^. These anti-PF4 antibodies are elicited through adenoviral vector associated adenoviral core protein in conjunction with somatic hypermutation with immunoglobulin light-chain allele IGLV3-21*02 or *03 found in selected human population ^18^. Independent from VITT, a low incidence of venous (VTE) and arterial (ATE) thrombotic events have also been reported from SARS-CoV-2 mRNA vaccination ^14^, implicating a second mechanism plus a direct role of SARS-CoV-2 spike protein to induce thrombotic events. Subsequent study further shows a direct correlation between elevated plasma SARS-CoV-2 spike protein and mRNA vaccination-associated myocarditis in a cohort of otherwise healthy young males ^19^.

One of the unique features of platelets is its surface-connected open canalicular system (OCS) that supports passive uptake of sub-micron size particles in platelets ^20,21^, a feature that is distinct from active phagocytic engulfment capability in neutrophils and macrophages. Although previous studies have shown virion particles can been seen with platelets using thin-section transmission electron microscopy (TEM) ^22,23^, it is unknown whether these virion particles are actively engulfed into platelets or passively trapped within OCS.

Published evidence has demonstrated that there are linkages between ABO blood types and COVID-19 infection (including risk of coronary artery disease) ^24,25^. It is perhaps less appreciated that the determinants of blood type are based on surface glycans, which modulate interaction with erythrocytes via sialoglycoconjugates ^26^. In contrast with the high-mannose enriched glycan shield in HIV surface proteins ^27^ that moonlights as molecular Velcro to facilitate infection ^28^, the SARS-CoV-2 spike proteins are enriched with sialoglycoconjugates across many of the 22 N-linked glycosylation sites per protein monomer ^29^. The high density of sialic acid termini of SARS-CoV-2 spike resembles the abundant sialoglycoconjugate on human Von Willebrand factor (VWF) ^30,31^ from its 12 N-glycan sites and 10 O-glycan sites per protein monomer. VWF acts as a haemostatic agent during platelet activation ^32^. Lowering the levels of VWF sialylated glycoconjugates represses VWF function and plasma half-life ^30^, but it is not clear whether sialic acids associated with theSARS-CoV-2 spike protein contribute to the pathogenetic functional impacts of SARS-CoV-2. A recent genome-wide association study (GWAS) has shown a nucleotide polymorphism of *β-galactoside α-2,6-sialyltransferase* (*ST6GAL1*), encoding an enzyme that adds sialic acid to glycoproteins, is strongly associated with SARS-CoV-2 infection, but the mechanistic contribution of *ST6GAL1* to SARS-CoV-2 infection and pathogenesis remains an enigma ^33^.

## Results

### SARS-CoV-2 spike protein-coated particles are trapped and remodelled invaginated platelet structures

As platelet activation has been linked to SARS-CoV-2 pathogenesis ^22,34^, we examined whether viruses could activate platelets to reorganise OCS structures (Fig. 1A-E). Quantitative ultrastructural analysis showed a significant increase with not only the size of platelets (Fig. 1C) but also the OCS-to-platelet area ratio, following SARS-CoV-2 spike coated particles activation (Fig. 1D). This activation was directly linked to an elongation of platelet OCS shape, with a 41.7 % and 26.8 % increase in both long- and short-axis, respectively (Fig. 1E), reducing platelet circularity from 0.34 to 0.26 (p<0.05, Fig. S1) as a consequence. Our data demonstrated that SARS-CoV-2 spike protein coated particles induced substantial membrane network remodelling that is consistent with cytoskeletal rearrangement and membrane extension.

**Figure 1.**
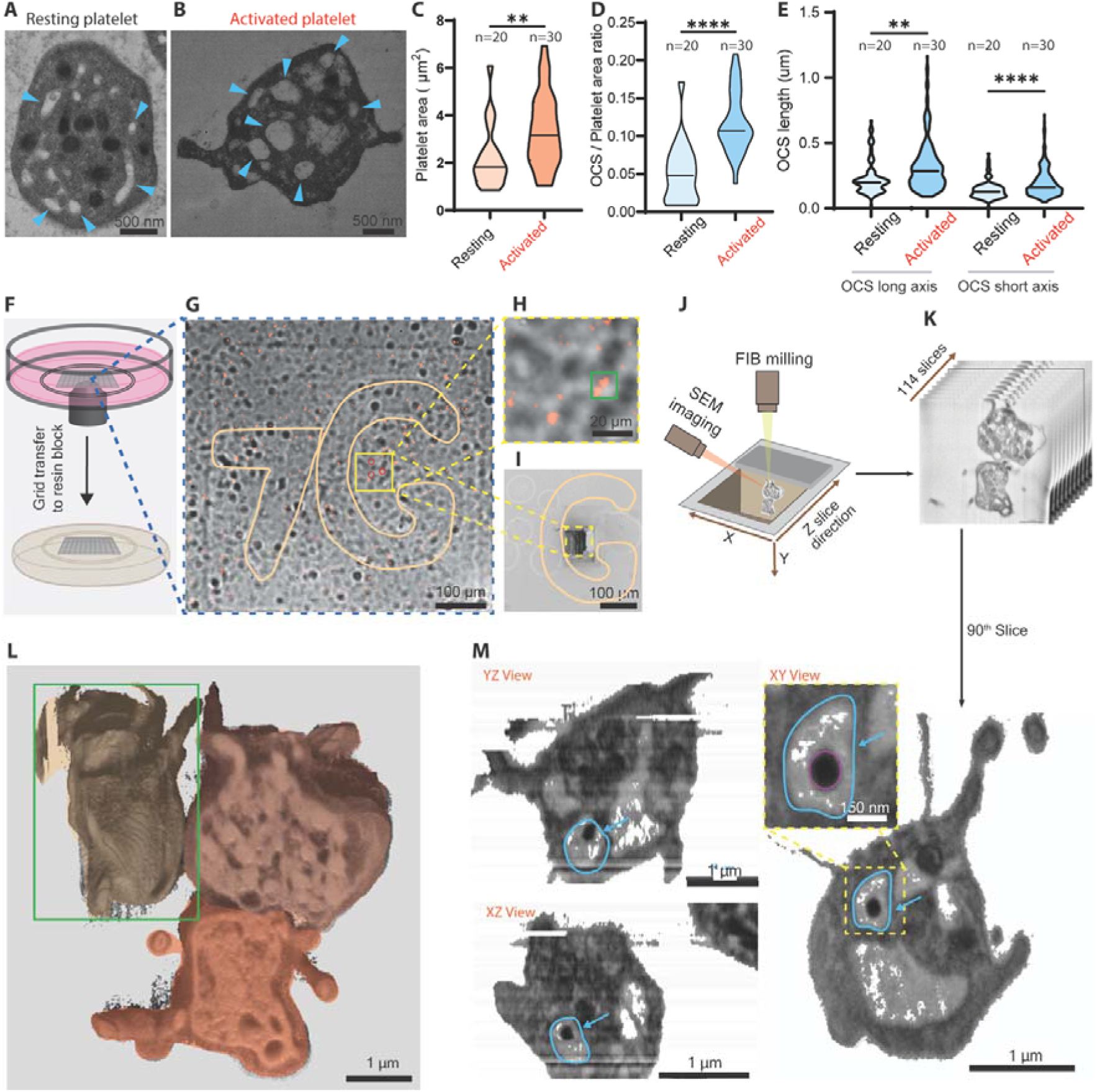
SARS-CoV-2 spike-protein coated particles induced remodelling of platelet ultrastructure and spike protein-coated particles were sequestered into the open canalicular system (OCS). Thin-section transmission electron microscopy (TEM) images of (A) resting-and (B) activated-platelets were shown, in which OCS were highlighted with blue arrows. Quantitative comparison showed different 2D areas of resting- (light orange) and activated-(dark orange) platelets (C, p<0.01). OCS area to platelet area ratio (D, p<0.0001) and OCS dimension comparisons (E, long- [p<0.01] and short-axis [p<0.0001]) between resting- (light blue) and activated (dark blue) platelet were shown. (F) Schematic diagram shows transfer of gridded coverslip alphanumeric coordinates from confocal imaging to the resin block for targeted EM. Correlation of (G-H) confocal microscopy image and (I) focused ion beam–scanning electron microscopy (FIB–SEM) images coordinates using the alphanumeric fiducial, ‘7G’ was highlighted for visualisation purposes. (H) is the zoom-in section with higher magnification for fluorescent image (targeted platelets highlighted in green square corresponding to L), while (I) was the same section post-FIB milling. (J) Serial FIB–SEM 20 nm slice image acquisition and (K) volumetric datasets spanning >100 sequential sections of platelet for volumetric ultrastructure analyses (L) 3D reconstruction of platelet aggregates. Green rectangle in (L) highlighted the section of platelet underwent the orthogonal slice view extraction in (M). (M) The orthogonal slice views from three vantage points (XY, XZ, YZ) confirm localisation of spike-coated particles within the three-dimensional architecture of the platelet OCS (highlighted with blue circumference and arrow).

Earlier studies showed that SARS-CoV-2 particles ^22^, as well as other viruses ^23,35^, can be identified within internal platelet structures using 2-dimensional thin section TEM, but it was unclear whether these particles were actively engulfed into internal vacuoles or passively captured via platelet OCSs. We employed a correlative light and electron microscopy (CLEM) workflow to track mCherry-tagged SARS-CoV-2 spike protein coated particles within platelets. A schematic overview of the experimental pipeline is shown in Fig. 1F-I. Platelets mixed with fluorescent-tagged virion particles were fixed onto an alphanumerically labelled EM gridded support (Fig. 1F). Spatial locations of platelets were first identified with low magnification confocal microscopy imaging (Fig. 1G), which was followed by selecting a few candidate virion-associated platelets (Fig. 1H). The chosen regions were subjected to focused ion beam scanning electron microscopy (FIB–SEM) milling (Fig. 1I) in accordance with an alphanumeric fiducial coordinates positioning. Serial FIB–SEM 20 nm think slice imaging was performed to collate a volumetric view of platelet ultrastructure (Fig. 1J), thereby generating three-dimensional (3D) datasets spanning >100 sequential sections covering over 2 μm depth in slicing direction (Fig. 1K-L). Reconstruction and segmentation of these datasets revealed the presence of viral particles trapped within the 3D space of platelet OCS. Orthogonal views (XY, XZ, YZ) extracted from the reconstructed volume (Fig. 1L) confirmed that these particles were localised within the 3D architecture of the invaginated OCS structures (Fig. 1M, Fig. S2-4 and Video S1). This volumetric validation provided direct spatial evidence of viral sequestration within an invaginated membrane platelet OCS system. Importantly, our 3D reconstruction of platelet aggregates demonstrated that viral particles were trapped within activated platelet aggregates (Fig. 1L). It has been suggested that SARS-CoV-2 spike protein engages with integrin for platelet activation ^36^. Luciferase reporter virus-bound platelet assay showed SARS-CoV-2 spike coated particles strongly associated with pre-chemically-fixed platelets upon mixing (Fig. S5). The ability of inactivated (chemically-fixed) platelets to capture particles is consistent with the passive covercyte mechanism ^20^ of OCS (Fig. S5). Trypsin-induced uncoupling of virion-platelet association illustrated viral particles were in a solvent accessible environment outside of the plasma membrane, including both the outer surface and the spatial cavities in OCS (Fig. S5). The ability of virion particles to bind with platelets reduces the sensitivity of virus detection in blood, particularly given blood plasma (platelet-depleted) is the gold standard in sampling blood-associated viral pathogens.

### SARS CoV-2 spike protein-coated particles drive platelet association and aggregation

Our 2D TEM data (Fig 1A-B) and 3D FIBS-SEM (Fig. 1K-M) showed specifics on the ultrastructural relationship between platelets and SARS-CoV-2 spike protein-coated particles. To evaluate the population-relationship between platelets and virion particles, confocal imaging showed that SARS-CoV-2 spike protein-coated particles substantially increased: (i) platelet-platelet interactions; and (ii) fluorescent-tagged virion–platelet association (Fig. 2A-B), when SARS-CoV-2 spike protein-coated particles were present. The significant increase of platelet-associated fluorescence intensity with SARS-CoV-2 spike protein-coated particles (Fig. 2C, p<0.05, N=6) suggested a role for the SARS-CoV-2 spike glycoprotein in the process. Qualitative differences in SEM determined platelet morphology were observed upon incubation with virion particles (Fig. 2D-H). Platelets incubated with SARS-CoV-2 spike protein-coated particles displayed visible aggregation (Fig. 2F-H), while untreated- (Fig. 2D) and naked (SARS-CoV-2 spike protein-free) particles treated- (Fig.2E) platelets remained largely dispersed. Platelets from both untreated- (Fig. 2D) and naked particles treated-(Fig.2E) samples exhibited minimal filopodia extension (Fig. 2D-E, Fig. S6). In contrast, platelets exposed to virion particles coated with either ancestral (Wuhan)- or Omicron- spike protein resulted in pronounced platelet aggregation (Fig. 2F-G), which were characterised by the presence of multiple platelets clustering and extensive nanoscale filopodia protrusions with heterogeneous morphology (Fig. 2H). To quantify the aggregation effects, platelet populations were classified based on the aggregated area sizes using SEM datasets from multiple donors (N = 9), incorporating 1825+ events per experimental condition (n=1825 to 2491). Platelet aggregates were binned into 4 size categories (i) 0–50 μm^2^; (ii) 50–100 μm^2^; (iii) 100–150 μm^2^; and (iv) >150 μm^2^ (Fig. 2K), equating to the joining of (i) 1-9; (ii) 9-17; (iii) 17-25, and (iv) >25 individual platelet ‘particles’, respectively. The majority (99.9%) of platelet population in both untreated- and naked particles-treated samples were confined to 50 μm^2^ (or <9 platelets), with the remaining 0.1% platelet aggregates assumed the size of 50–100 μm^2^ (Fig. 2K). In contrast, treatments of platelets with either ancestral- or Omicron-spike protein-coated particles led to the presence of large platelet aggregates exceeding 100 μm^2^ (Fig. 2K), which accounted for 0.1% or more of the platelet populations. The potential of Omicron spike protein-coated particles as a stronger inducer of platelet aggregates over ancestral (Wuhan) spike protein-coated particles requires further analysis (Fig. 2K). The ‘average size of platelet aggregates’ and the ‘fractional area occupancy of platelet aggregates within the platelet population’ were analysed by pooling samples derived from 9 donors, in which ∼4650 platelets aggregates were detected in the platelet population across all conditions (Fig. 2L-M). We defined platelet aggregate as a connected area consisting of two or more platelets. Our data showed that the mean platelet aggregate area significantly increased from 9 / 11 μm^2^ (for no- / naked-particles- treatment, Fig. 2L) to 13.5 / 14.5 μm^2^ (ancestral- / Omicro-spike protein-coated particles treatment, Fig. 2L, p<0.01). No mean area difference was observed between no- and naked-particles- treated platelet aggregates (Fig. 2L). There were over a 60% increase of ‘fractional areas occupied by platelet aggregates within the platelet population’ upon treatments with ancestral- (Fig. 2M, from 40% to 67%, p<0.01) or Omicron- (Fig. 2M, from 40% to 65% p<0.05) spike protein-coated particles. These data showed that virion-associated SARS-CoV-2 spike proteins activated platelets by increasing both the size and the frequency of platelet aggregation (Fig. 2K-M), in which the SARS-CoV-2 spike protein was a major contributor to platelet activation.

**Figure 2.**
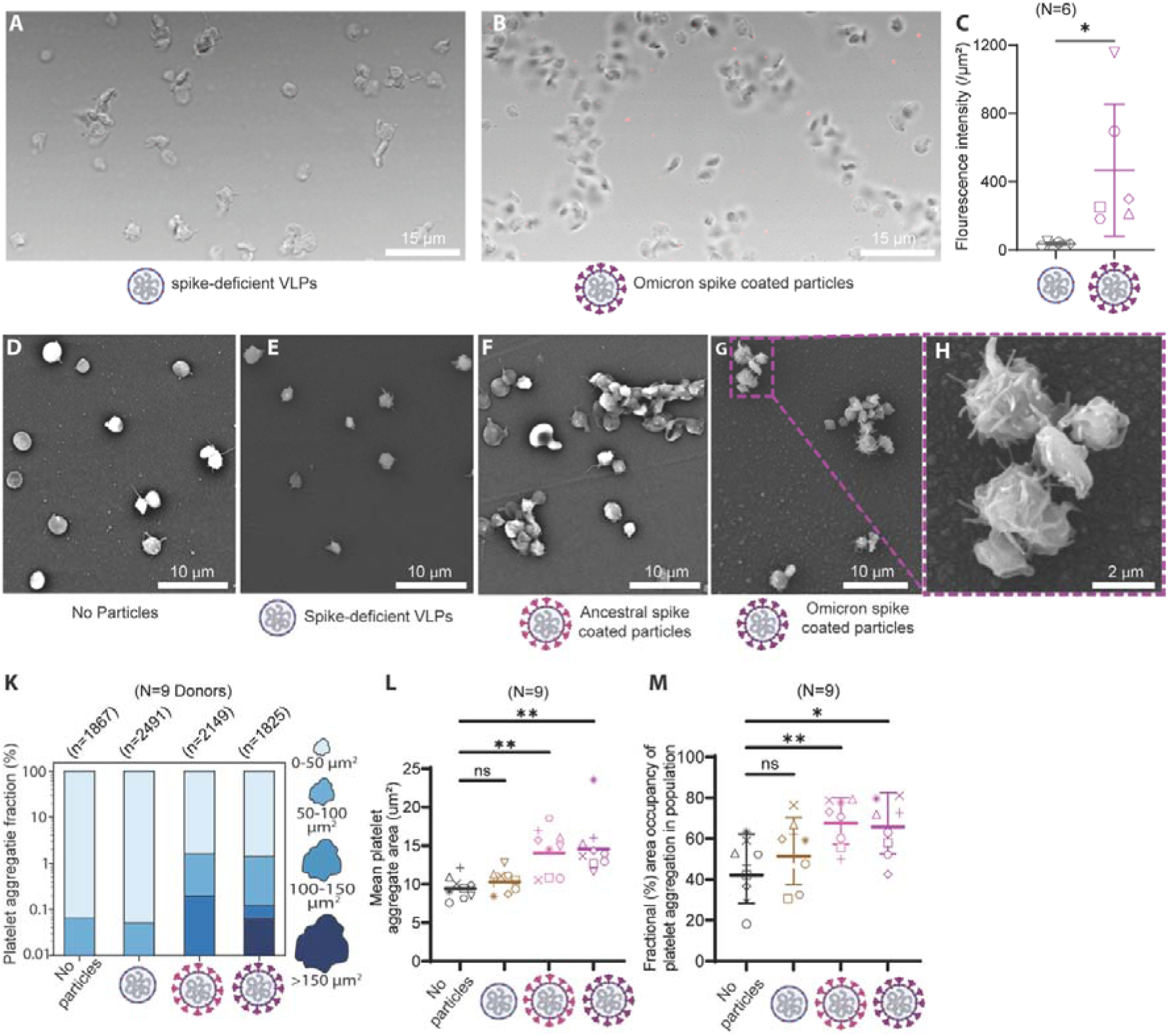
SARS-CoV-2 spike protein-coated particles promote platelet–platelet interaction and aggregation. Confocal imaging shows association of mCherry-labelled virion particles with platelets following incubation with (A) spike protein-deficient particles and (B) Omicron spike protein-coated particles. (C) Quantification of fluorescence intensity associated with platelets in the presence of spike protein-deficient and spike protein-coated particles (N=6). (D) SEM image of platelets (no particle), (E) SEM image of platelets incubated with spike protein-deficient particles (F) SEM image of platelets incubated with ancestral spike protein-coated particles (F) SEM image of platelets incubated with omicron spike coated particles (H) Higher resolution SEM images showing morphology of aggregated platelets and filipodia (K) Size-distribution analysis of platelet aggregates across donors (N = 9; >1800 events per condition) (L) Statistical comparison of mean aggregate area and (M) statistical comparison of fractional area occupancy of platelet aggregates within the platelet population. Capital ‘N’ is for Donor numbers and little ‘n’ is for number of events. Scale bars sizes are listed. *p<0.05, **p<0.01, ***p<0.001, ****p<0.0001.

### SARS CoV-2 spike protein alone induces dose-dependent platelet aggregation

To determine whether the spike protein alone was sufficient to drive platelet activation, platelets were exposed to increasing concentrations (from 0.25 μg/mL to 25μg/mL) of purified recombinant SARS-CoV-2 spike protein for morphological analyses with SEM. Data collected from platelets (across from 8 donors) consistently showed dose dependent recombinant spike protein-induced platelet aggregations (Fig.3A), illustrating the specificity of SARS-CoV-2 spike protein to promote platelet activations. The observed recombinant spike protein-induced morphological changes in platelets (Fig.3A, S6) were indistinguishable from those seen with platelet treatment using intact SARS-CoV-2 spike protein-coated particles (Fig. 2D-G). These observations suggest that the SARS-CoV-2 associated platelet aggregation capacity was primarily driven by the viral spike protein. Population analyses showed 0.25 μg/mL of recombinant SARS-CoV-2 spike was sufficient to increase the amount of 50-100 μm^2^ size platelet aggregates 5-fold (from 0.04% to 0.8%, Fig. 3B), while up to 25-fold increase of platelet aggregates (or 2% of the platelet population) exceeding 50 μm^2^ were detected with 25 μg/mL recombinant spike protein stimulation (Fig. 3B). Roughly 0.1% of platelets reached 150 μm^2^ or larger in size upon 25 μg/mL recombinant spike protein treatment (Fig. 3B), which was associated with an increase to 15 μm^2^ (p<0.01) mean platelet aggregate area from 8 μm^2^ in the untreated control (Fig. 3C). The fractional area occupancy of platelet aggregate within the platelet population had an increase from 50% in untreated control to roughly 65%, 70%, and 75% (Fig. 3D) upon treatments with 0.25 μg/mL, 2.5 μg/mL and 25 μg/mL, respectively, of the recombinant spike proteins (Fig. 3D). Our results demonstrated that platelet activation and aggregation were driven by direct molecular interactions with the SARS-CoV-2 spike protein that was independent from intact viral particles. Our findings are consistent with previous studies showing SARS-CoV-2 can directly interact with and activate platelets, thereby contributing to thrombo-inflammatory responses observed in COVID-19 ^6,22,34^. Furthermore, our observation provided a scientific rationale to account for the molecular mechanism on how otherwise healthy male subjects experienced myocarditis from mRNA-based COVID vaccination in the absence of SARS-CoV-2 infection ^19^.

**Figure 3.**
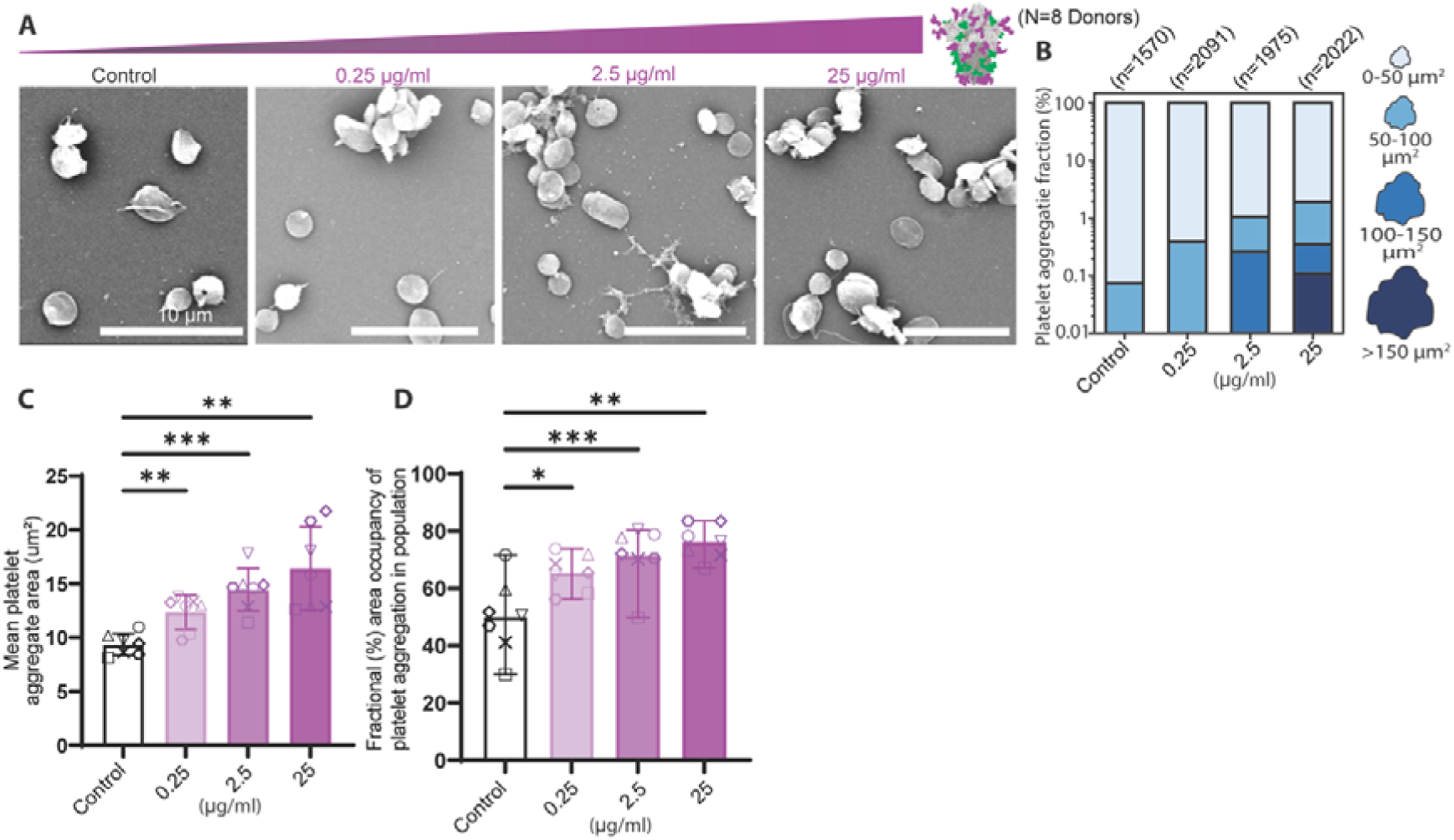
SARS-CoV-2 spike protein alone induces dose-dependent platelet aggregation. (A) Representative SEM images of platelets under control conditions and following incubation with increasing concentrations of purified ancestral SARS-CoV-2 spike protein (0.25, 2.5, 25 μg/mL). (B) Size distribution of platelet aggregates classified into bins (0–50, 50–100, 100–150, >150 μm²). Statistical analysis of (C) mean aggregate area and (D) fractional area occupancy of platelet aggregate within the platelet population. Capital ‘N’ is for Donor numbers and little ‘n’ is for number of events. Scale bars sizes are listed. *p<0.05, **p<0.01, ***p<0.001, ****p<0.0001.

### Spike protein terminal sialylation drives platelet aggregation

N-linked glycosylation of proteins can be manipulated by inactivation of host cell glycosylation enzymes during protein synthesis or exogenous glycosidases in purified recombinant proteins (See schematic in Fig. S7). To identify molecular determinants within the SARS-CoV-2 spike proteins that were responsible for platelet activation, we leveraged the N-acetylglucosaminyl transferase I defective (GnTIL) cells line to produce glycosylation-modified SARS-CoV-2 spike protein coated-particles ^28^. The SARS-CoV-2 spike protein produced in GnTILcells contained high-mannose N-linked glycans across all 66 N-glycan sites per each of the SARS-CoV-2 trimer (Fig. 4A, S7), in complete contrast to the sialic acid-rich complex glycans found on authentic SARS-CoV-2 spike protein produced in the control cell line (Fig. 4B). Platelets incubated with Omicron spike protein-coated particles containing high mannose glycans remained largely as individual ‘units’ with minimal aggregation (Fig. 4A). In contrast, platelets exposed to sialic acid-enriched spike protein-coated particles formed many platelet-platelet interactions that were associated with elevated levels of fluorescent signal (Fig. 4B). Quantitation of particle-associated fluorescence across platelets from 6 donors between two types of SARS-CoV-2 glycan spike protein-coated particles were not significance (Fig. 4C). Similarly, employing an *in vitro* luciferase-reporter virus-platelet binding assay with platelets derived from 3 donors did not show a significant difference in the capacity of these two types of SARS-CoV-2 spike protein glycans to interact with platelets. The qualitative differences platelet clustering between these two types of glycan-decorated SARS-CoV-2 spike protein-coated particle suggests a role of glycans in platelet activation (Fig. 4A-B). SEM analyses showed platelets incubated with high-mannose SARS-CoV-2 spike protein-coated particles exhibited background to low-level of detectable activation (Fig. 4E, S8) in comparison to a no particle incubation control (Fig. 4D, S8). In contrast, sialic acid-containing glycosylated SARS-CoV-2 spike protein-coated particles promoted platelet activation resulting in more frequent nano-size filopodia protrusions (Fig. 4F). Population analyses on the sizes of platelet aggregates confirmed high-mannose SARS-CoV-2 spike protein-coated particles were not effective in activating platelets *in vitro* (Fig. 4G, N=9, n=1898), exhibiting either no changes in mean platelet aggregate area from control at ∼10 μm^2^ (Fig. 4H) or non-significant differences with 40-50% fractional area occupancy of platelet aggregate (Fig. 4I). The distinct impacts between (i) high mannose- and (ii) complex sialylated glycans-, SARS-CoV-2 spike protein-coated particles on platelet aggregation (Fig. 4D-I) suggested the precise composition of terminal glycans in the SARS-CoV-2 spike glycosylation could be a determinant in platelet clotting induction and SARS-CoV-2 associated thrombosis.

**Figure 4.**
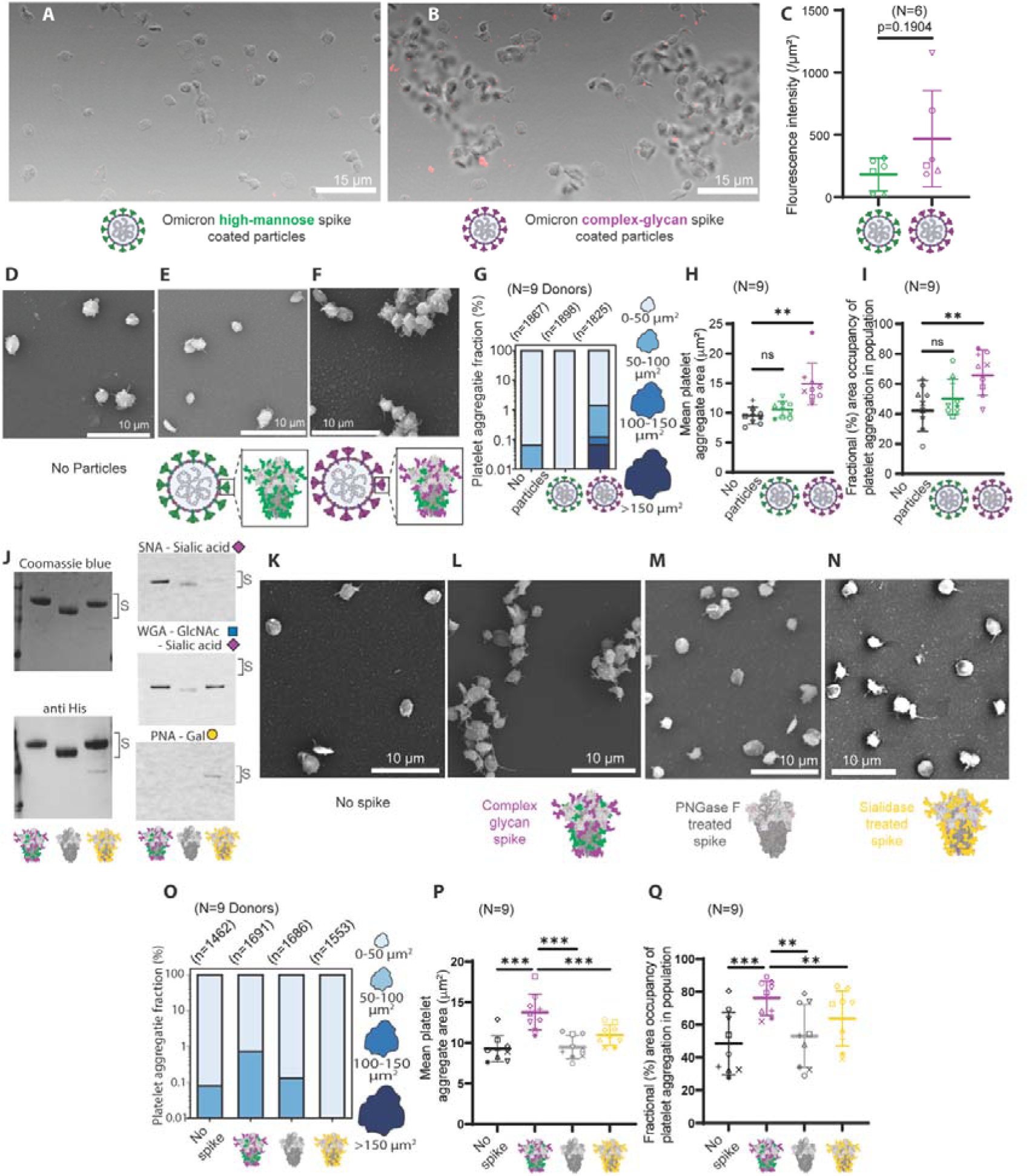
SARS-CoV-2 spike protein terminal sialic acid residues induce platelet aggregation. (A-B) Confocal images of platelets incubated with mCherry-labeled Omicron spike protein-coated particles possessing (A) high-mannose (GnTI-/- HEK293T) or (B) complex-type (HEK293T) glycans. (C) Quantification of VLP-platelet association with particles with high mannose or particles with complex glycan. (D-F) Scanning electron microscopy (SEM) images of human platelets that have been incubated with: (D) no particles, (E) Omicron spike high-mannose glycoprotein-coated particles, and (F) Omicron spike complex glycoproteins-coated particles. (G) Quantification of mean platelet aggregate areas (G) and platelet aggregation index (H) from SARS-CoV-2 spike protein-coated particles induced platelet aggregation SEM analyses. (I) Size distributions of platelet aggregates from D-F. (J-M) SEM images of human platelets that have been incubated with different recombinant spike proteins: (J) Recombinant protein blots visualised by Coomassie blue, anti-his antibody, or lectins (*sambucus nigra* lectin [SNA], wheat germ agglutinin [WGA], and peanut agglutin [PNA]) for the detection of recombinant spike protein and associated glycans. (K) no spike protein; (L) complex glycan spike protein; (M) glycan diminished PNGaseF treated spike protein; (N) sialic acid diminished sialidase-treated spike protein. (O) Size distributions of platelet aggregates from K-N. Quantification of mean platelet aggregate areas (P) and fractional area occupancy of platelet aggregate in platelet population (Q) from recombinant SARS-CoV-2 spike protein induced platelet aggregation SEM analyses (K-N). Capital ‘N’ is for Donor numbers and little ‘n’ is for number of events. Scale bars sizes are listed. *p<0.05, **p<0.01, ***p<0.001, ****p<0.0001.

To dissect the molecular contribution of SARS-CoV-2 spike protein glycans to platelet aggregation and thrombosis, specific glycosidases were used to trim off selected groups of glycans to assess their capacity to induce platelet aggregation. Peptide:N-glycosidase F (PNGase F) was used to remove SARS-CoV-2 spike protein N-linked glycans at the base of the N-linked glycan ‘tree’ resulting in the complete removal of N-glycans (Fig. S7). A sialidase (α2-3,6,8,9 Neuraminidase A) was employed to snip off the terminal sialic acid residues on the N-linked glycans (Fig. S7), thereby maintaining the majority of N-glycan tree, while exposing galactose from previously sialic acid-capped structures to be the new terminal N-glycans structures. Polyacrylamide gel electrophoresis (PAGE) protein separation coupled with protein staining, immunoblot, and lectin probing were then carried out to confirm that: (i) the majority removal of N-glycans by PNGase F; and (ii) the terminal sialic acids were cleaved off from the N-glycans. These glycan modifications resulted in: (a) a faster migration of recombinant spike protein upon PNGase F treatment; (b) a lower detectable level of sialic acids from both PNGase F and sialidase treatment; and (c) an enhanced detectability of galactose terminal residues post-sialidase treatment (Fig. 4J).

Using (i) no-spike protein stimulated platelets and (ii) wild-type sialic acid-containing glycosylated SARS-CoV-2 spike protein stimulated platelets as negative and positive controls, respectively, in SEM analyses (Fig. 4K-L, S9), it was noted that the ultrastructural morphologies of platelets upon stimulation with either PNGase F-treated- (Fig. 4M, S9) or sialidase-treated- (Fig. 4N, S9) SARS-CoV-2 spike protein were closer aligned with the no-spike protein stimulated platelets negative control. PNGase F-treated-spike protein stimulated platelets (Fig. 4M, S9) had limited signs of activation in comparison with the sialic acid-containing complex glycan SARS-CoV-2 spike protein treated positive control (Fig. 4L, S9). Population analyses on the sizes of platelet aggregates showed N-glycan-depleted recombinant SARS-CoV-2 spike have only 0.1% platelet aggregates that are greater than 50 μm^2^ in size (Fig. 4O), which was at the background levels observed in no-spike protein stimulated platelets (Fig. 4O). Both ‘mean platelet aggregate area’ (Fig. 4P) and ‘fractional area occupancy of platelet aggregate’ (Fig. 4Q) from N-glycan-depleted recombinant SARS-CoV-2 spike protein treated platelets were compatible with background controls but significantly different from wild-type complex glycan SARS-CoV-2 spike protein stimulated platelets (p<0.001 Fig. 4P, p<0.01 Fig. 4Q). These data showed that the N-glycans of the SARS-CoV-2 spike protein were directly involved in platelet activation and aggregation.

More strikingly, precision removal of terminal sialic acid residues from the SARS-CoV-2 spike protein N-glycans resulted in an absence of platelet aggregates above 50 μm^2^ (Fig. 4O) and less visible nanoscale filopodia extrusion (Fig. S9). Both ‘mean platelet aggregate area’ (Fig. 4P) and ‘fractional area occupancy of platelet aggregate’ (Fig. 4Q) from sialic acid free (or galactose terminal-capped) spike protein-stimulated platelets were significantly different from the wild-type spike protein-induced platelet control (p<0.001 Fig. 4P, p<0.01 Fig. 4Q). These data provided direct evidence that the terminal sialic acid residues on the SARS-CoV-2 spike protein were responsible for platelets aggregations *in vitro*, suggesting a role for SARS-CoV-2 spike protein’s sialic acid residues in COVID-19 associated thrombo-inflammatory pathology.

## Discussion

Platelet hyperactivation and thrombotic events are hallmarks of severe SARS-CoV-2 infection, yet the molecular determinants underlying these processes remain incompletely defined. Our data showed that SARS-CoV-2 spike protein-coated particles and the spike protein directly induced platelet aggregation. We further identified spike protein glycosylation as a critical regulator of this effect. Using imaging and quantitative analyses, we showed that spike protein-coated particles promote platelet aggregation, and spike protein alone was sufficient to cause platelet aggregation in a concentration-dependent manner. Importantly, using glycan modified SARS-CoV-2 spike protein-coated particles and glycosidase trimmed recombinant SARS-CoV-2 spike protein, we directly demonstrated that the terminal sialic acid residues of SARS-CoV-2 spike protein are the key determinant which results in platelet aggregation induction. Together, these findings establish a mechanistic link between SARS-CoV-2 spike protein-associated sialic acid residues and platelet aggregation thrombosis biology.

A recent study has shown that platelet OCS can trap cell-free DNA ^21^. Diagnostic detection of rare foetal DNA or cancer-derived DNA is more sensitive with lysed whole blood in comparison with ‘conventional gold standard’ platelet-depleted plasma ^21^. The ability of platelets to trap SARS-CoV-2 spike protein-coated particles in OCS is likely to: (i) impair blood detection of SARS-CoV-2 using standard platelet-depleted plasma; and (ii) enable platelets to serve as a Trojan horse reservoir to delay virus clearance in convalescence individuals, thereby prolong the duration of SARS-CoV-2 infection in the host.

It is estimated that 14.4 million life-years have been saved with the SARS-CoV-2 vaccine ^37^, yet the unappreciated biology that contributes to SARS-CoV-2 related thrombosis ^1,2,4^ and vaccine-associated thrombotic events ^14,16,38,39^ have dampened vaccine confidence. While the mechanism of pathogenetic anti-PF4 antibodies induction in VITT is now resolved ^18^, the source of SARS-CoV-2 mRNA vaccine-related thrombotic events ^14^ and myocarditis ^19^ remain a mystery. Building from our prior work on glycan-glycan interaction in viral pathology ^28^, we noted that the high density of terminal sialic acid residues is found on the surfaces of both the SARS-CoV-2 spike proteins ^29^ and the host haemostasis initiator Von Willebrand factor (VWF) ^30,31^. As N-glycan-associated terminal sialic acid residues contribute to VWF functions and prevent premature plasma clearance ^40–42^, we postulated this shared feature enables the SARS-CoV-2 spike protein to mimic host cell VWF to activate platelets that leads to the formation of microscopic scale clots resulting in thrombotic thrombocytopenia.

Our proposed mechanism on viral protein-associated sialic acids as the causation of SARS-CoV-2 induced thrombotic thrombocytopenia is in line with recent work showing that a sialyltransferase gene *ST6GAL1* is associated with SARS-CoV-2 infection ^33^. The contribution of *ST6GAL1* intron single nucleotide polymorphism (SNP) to SARS-CoV-2 infection could be through: (i) altering the stability of *ST6GAL1* mRNA for protein expression; and/or (ii) interfering with the functions of other glycosylation machinery ^33^. With up to 2% of the human genome (over 400 glyco-genes plus 950 pathway-process related genes) directed towards glyco-modifications, personalised glycosylation does not only determine human blood types ^43^ but also dictates the individualised glycan profiles of the SARS-CoV-2 spike protein upon infection. If the patterns of SARS-CoV-2 spike protein terminal sialic acid residues and their relationships with host biology are the keys to initiate COVID-19 thrombotic pathogenesis, it is perhaps not surprising to observe the vast differences on the level of disease pathogenesis across SARS-CoV-2 infected individuals.

Despite the high-level safety standard being achieved with the adenoviral vector-based SARS-CoV-2 vaccine ^11^, the concerns with rare vaccine-induced thrombotic thrombocytopenia have forced several effective COVID vaccines out of the market, prematurely. Identified the linkage between adenoviral core proteins and immunoglobulin light-chain allele, IGLV3-21*02 or *03 will help mitigate risks of CVDs through adenoviral vector vaccine-related VITT ^18^ but does not offset rare SARS-CoV-2 mRNA vaccine-associated thrombotic events ^14^ or myocarditis ^19^. As we have shown that glycan-modified SARS-CoV-2 spike protein is ineffective in platelet aggregation induction, glycan-modified vaccines have the potential to reduce risks of vaccine-related thrombotic events. Recent studies have shown that introduction of glycosylation machinery inhibitors during viral protein production can disable SARS-CoV-2 function by interfering with the glycan profiles of progeny viruses ^44^. Local co-injection of glycosylation machinery inhibitors with SARS-CoV-2 mRNA vaccine could alter the glycan profiles of soluble SARS-CoV-2 spike protein to reduce the risk of SARS-CoV-2 mRNA vaccine-associated thrombotic events or myocarditis, while still enabling the production of the protein antigen to elicit protective immunity. Such a ‘glycan-modified mRNA vaccine’ approach is likely to have wider applications to other glycan-dependent pathogens, including the potential to crack open the HIV glycan shield and to expose broadly neutralising epitopes, thereby facilitating the elicitation and the maturation of protective immunity against HIV.

## Materials and Methods

### Human Ethics statement

Human blood samples were obtained from healthy adult donors following informed consent in accordance with the Declaration of Helsinki. Ethical approval for this study was granted by the Griffith University Human Research Ethics Committee (HREC approval number: HREC/2026/0129). All participants provided written informed consent prior to blood collection, and all experiments involving human blood were conducted in accordance with approved institutional guidelines.

### Blood collection and platelet isolation

Human blood was collected in Acid Citrate Dextrose (ACD) solution containing BD Vacutainer® glass whole blood ’ACD-A’ tube from the de-identified donors. Platelet-rich plasma (PRP) was prepared from whole blood collected from healthy adult donors. All donors were medication-free for at least 10 days prior to blood collection and had not taken antiplatelet or anti-inflammatory drugs during this period. Blood was collected using standard venipuncture techniques with minimal stasis to avoid artifactual platelet activation. To minimise spontaneous activation, PRP was prepared under controlled conditions using low-speed centrifugation without brake, and all handling steps were performed gently using wide-bore pipette tips. Samples were processed at room temperature and used within a standardised time frame following collection.

Immediately after blood was drawn from the donors, it was centrifuged for 15Lmin at 100L×L*g* at room temperature. The top 2/3 of the PRP was transferred to a fresh Falcon tube. After gently inverting 3-5 times to homogenise the concentration and distribute to 1.5 ml tubes each with 95 ul. Immediately, spike protein was added to the PRP with no pipetting or swirling the pipette tip. After incubating the PRP and spike protein mix at room temperature or 37 °C for 15-30 minutes, samples for SEM imaging were fixed with 2.5% glutaraldehyde (end concentration)

### Production of SARS-CoV-2 spike-coated- and naked- virus-like particles (VLPs)

SARS-CoV-2 spike-coated- and naked (spike-free)- virus-like particles (VLPs) were produced using the HEK 293T cell line or GlcNAc transferase I defective 293S GnTI-/- cells via transfection. All cells were cultured under standard conditions as previously described ^28,45^. The plasmids used to generate the nanoluciferase VLPs for the cell surface binding assay were: (i) NL4-3 based proviral DNA construct expressing Gag and Env, but with deletions in RT, IN, Vif and Vpr (pNL4-3 ΔRTΔINΔVifΔVpr); and (ii) nanoluciferase-Vpr expression construct under control of the CMV promoter. Each transfection utilised a total of 12 μg plasmid DNA, combined in an 8:1 ratio of each plasmid listed above. The plasmids used to generate the VLPs for fluorescent imaging were: (i) pNL43ΔPolΔEnv; (ii) pNL43ΔPolΔEnv-Gag-mEOS2; and (iii) SARS-CoV-2 spike protein expression vector. Each transfection utilised a total of 12 μg plasmid DNA, combined in a 3:1:1 ratio of each plasmid listed above. For VLP production, HEK 293T cells were transiently transfected using polyethylenimine (PEI MAX). PEI/DNA complexes were formed by adding PEI to plasmid DNA diluted in a small volume of fresh DMEM. The mixtures were vortexed vigorously and incubated at room temperature for 30 min prior to transfection. Culture supernatants of transfected HEK 293T cells were harvested 48 h post transfection and clarified by centrifugation at 1500 x g for 10 min. Cell debris was removed by passing the supernatants through a 0.45 μm filter. VLPs were then concentrated by ultra-centrifugation over a 20% sucrose cushion. Concentrated VLPs were resuspended in DPBS and stored at −80 °C. VLP production, where possible, was quantified using a p24 ELISA (Xpress Bio).

### Recombinant spike protein expression and purification

Plasmid encoding for Hexaprolin spike FL trimer was transiently transfected into HEK293-T cells at a final concentration of 3 µg/mL in Freestyle™ 293 Expression Medium using polyethylenimine (PEI, Polysciences®) as a transfection reagent. After 24 h, the cells were diluted 1:1 with ExCell® 293 Serum-Free Medium for HEK 293 cells supplemented with 2.2 mM valproic acid to promote additional endocytic uptake of the plasmid-PEI complex. After 4 days, the supernatant was clarified by centrifugation for 30 min at 3000 × g at 4°C and treated with benzonase® (Merck) for 4 h at 4°C before downstream purification.

A pre-packed Ni^2+^-sepharose column (HisTrap Excel, GE Life Sciences) was used to purify His-tagged proteins from the cell culture supernatant by immobilised metal ion affinity chromatography (IMAC). The Ni^2+^-column was first equilibrated with 50 mM Na_2_HPO_4_, 300 mM NaCl, and 10 mM Imidazole pH 8 for capturing of the His-tagged Hexaprolin spike FL protein. Following, the protein was eluted with 250 mM imidazole in the same buffer and detected by UV absorbance at 280 nm. TEV protease cleavage was achieved before the protein was concentrated and buffer exchanged into 20 mM Tris and 200 mM NaCl (pH 8.0) over a 30 KDa cut-off Amicon Centrifuge Filter. The concentrate was loaded on a HiLoad 26/600 Superdex pg 200 column (GE Life Sciences) for size exclusion chromatography and collected in fractions of 1 mL. The fractions were checked for protein presence and purity by SDS-PAGE with colloidal Coomassie blue staining to visualise protein bands After expression and purification, Hexprolin spike FL protein was concentrated to 15 mg/mL and stored at – 20 °C until further use.

### SEM imaging and image analysis

For scanning electron microscopy (SEM) imaging, platelet-rich plasma (PRP) samples (95 µL) were incubated with either spike-coated particles (5 µL; 40 ng/µL final particle stock concentration) or soluble spike protein for 15–30 minutes at room temperature or 37 °C. For dose-dependent experiments, spike protein was added at final concentrations of 0.25 μg/mL, 2.5 μg/mL, and 25 μg/mL in PRP. In all other experiments, spike protein was used at a final concentration of 25 μg/mL. Control samples received 5 µL of gel filtration buffer, matching the buffer used for spike protein preparation. Immediately after incubation, samples were fixed by adding 2.5% glutaraldehyde to reach a final volume of approximately 400–500 µL and fixed for 30 minutes. This dilution step with glutaraldehyde was critical, as fixation without dilution caused PRP samples to become gel-like; maintaining a liquid suspension ensured efficient platelet settling onto substrates. Following 30 minutes fixation, platelet suspensions were deposited onto poly-L-lysine–coated Thermanox coverslips and allowed to settle and adhere to the substrate and continue fixing overnight. Samples were washed twice with PBS anddehydrated with serial increasing concentrations of ethanol (30%-50%-70%-90%-100%) for 10 minutes at each ethanol concentration, and twice with 100% ethanol. The samples were then dehydrated using a critical point dryer (Leica EM CPD300). The dried samples on Thermanox substrates were mounted onto aluminium specimen stubs using conductive carbon tape, and the edges of the substrates were additionally covered with carbon tape to improve electrical conductivity. Samples were subsequently sputter-coated with gold prior to imaging.

SEM imaging was performed under identical, standardised acquisition conditions using a Thermo Fisher Phenom XL SEM. At least six random, non-overlapping fields were captured per donor for each experimental condition. Following image acquisition, all SEM images were analysed using ImageJ. A platelet aggregate was defined as the association of any two or more platelets (total area > 6 µm²). The surface areas of individual platelets and platelet aggregates were quantified using ImageJ, and at least 6 images (technical replicates) platelets area was averaged per donor prior to statistical analysis.

### Thin-section TEM imaging of platelets

Platelet-rich plasma (PRP) was prepared from whole blood by 15 minutes of low-speed centrifugation (100 × g). Platelet activation was induced by adding 5 µL of Omicron spike–coated particles (80 ng/µL) to 95 µL of PRP, followed by incubation at 37 °C for 30 minutes. Resting platelet controls were prepared by incubating PRP under identical conditions without the addition of any agonist for 30 minutes at 37 °C. Following incubation, samples were fixed overnight at 4 °C in 2.5% glutaraldehyde prepared in PBS. During fixation, PRP samples were diluted approximately 3–5-fold with fixative to ensure proper preservation of platelets as in liquid suspension. Samples were washed twice with PBS and followed by 1% osmium tetroxide post-fixation using a Pelco Biowave processor following our methods reported elsewhere ^46^. Samples were dehydrated through a graded ethanol series (30–100%) and infiltrated with LX112 resin, followed by polymerisation at 60 °C for 48 h. Resin blocks were trimmed and sectioned (∼70 nm thickness) using a Leica Ultracut UC7 ultramicrotome with a DiATOME diamond knife. Sections were collected onto Formvar-coated TEM grids, stained with 5% uranyl acetate and Reynolds’ lead citrate, air-dried, and imaged using either a JEOL JEM-1400 Flash or Hitachi HT7700 operating at 80 kV. Thin-section TEM images of platelets and their OCSs parameters were analysed using the ImageJ software.

### Focus Ion Beam – Scanning Electron Microscopy (FIB-SEM) analysis

mCherry-tagged Omicron spike–coated particles were incubated with platelet-rich plasma (PRP) for 30 min at room temperature or 37 °C and fixed in 4% paraformaldehyde (PFA). PFA fixed samples were transferred onto gridded glass-bottom dishes (MatTek gridded glass-bottom dish) to enable coordinate-based correlation between fluorescence and electron microscopy images. Confocal fluorescence imaging was performed to identify regions of interest (ROIs). Following confocal imaging, samples were post-fixed in 2.5% glutaraldehyde and processed using a workflow like that used for thin-section transmission electron microscopy sample preparation in this work. To enhance membrane contrast for focused ion beam scanning electron microscopy (FIB–SEM), samples were further post-fixed in 1% osmium tetroxide overnight at 4 °C, washed twice with PBS, and subjected to an additional osmium tetroxide staining step using a Pelco BioWave microwave processor.

Samples were then embedded in LX112 resin and polymerised at 60 °C for 24–48 hours. After resin curing, the plastic edges surrounding the glass coverslip were trimmed, and the resin block was released from the coverslip by alternating cycles of liquid nitrogen and hot water (2–5 cycles), yielding a grid-transferred resin block approximately 3–5 mm thick. Prior to imaging, the resin block surface was sputter-coated with platinum (5-10 nm) and transferred to a FIB-SEM. Low-magnification SEM overview images were acquired to identify alphanumeric grid coordinates corresponding to previously imaged ROIs from confocal microscopy. After reidentifying of the ROI in FIB-SEM, protective carbon and platinum layers were deposited over the block surface to minimise ion-beam damage during milling. Initial coarse trench milling was performed manually. Once platelet structures were exposed, automated serial slice-and-view acquisition was initiated. Three-dimensional datasets were collected at a voxel size of 5.4 × 5.4 × 20 nm with a slicing thickness of 20 nm.

### Luciferase reporter virus-like particles (VLPs) – Platelets binding assay

NanoLuc-tagged virus-like particles (VLPs) pseudotyped with Omicron spike or spike-deficient control were generated in HEK293T cells or GnTi- cells and quantified by p24 ELISA prior to use. Platelet-rich plasma (PRP) was prepared from citrated whole blood by centrifugation at 100 × g for 15 min at room temperature without brake, and platelets were maintained under resting conditions throughout preparation. Washed platelets (1 × 10L per condition) were incubated with NanoLuc-tagged VLPs at defined particle-to-platelet ratios (approximately 100:1) in a final volume of 200 µL at 37 °C for 30 min with gentle mixing. Following 30 minutes incubation at 37 °C, platelets were pelleted at 1200 × g for 10 min and washed five times with Tyrode’s buffer to remove unbound particles. Platelet pellets were then divided into three fractions to quantify total associated, surface-bound, and internalised VLPs. Total platelet-associated NanoLuc signal was measured directly after resuspension in buffer and addition of substrate. Surface-bound and partially internalised VLPs were removed by brief treatment with 0.05% trypsin at 37 °C for 3–5 minutes followed by washing 3 times.

### PNGase F and Sialidase treatment of spike protein under non-denaturing conditions

Recombinant His-tagged SARS-CoV-2 spike protein was diluted to 0.556 mg mLL¹ prior to enzymatic treatment. Deglycosylation reactions were performed using PNGase F (for the removal of N-linked glycans) and sialidase (for the removal of terminal sialic acid residues), and the mock (without enzyme treatment) respectively. All reactions were incubated at 37 °C for 1 h. Following the manufacturer’s recommended procedure, for PNGase F treatment, 100 µg spike protein (180 µL) was mixed with 20 µL 10× GlycoBuffer and 30 µL PNGase F enzyme. For sialidase treatment, 30 µg spike protein (54 µL) was mixed with 6 µL 10× GlycoBuffer and 30 µL sialidase enzyme. Mock-treated spike controls were prepared by incubating spike protein with 10× GlycoBuffer only under identical conditions. Following incubation, reaction mixtures were subjected to buffer exchange and enzyme removal using 100 kDa molecular weight cut-off Amicon centrifugal filters (0.5 mL). Samples were centrifuged at 5000 ×g for 5 min at 4 °C, followed by replenishment with gel-filtration buffer to the retained volume. This wash step was repeated five times to ensure the removal of free enzyme and reaction buffer components. Protein concentrations following deglycosylation and buffer exchange were determined by absorbance at 280 nm using a Nanodrop spectrophotometer and the concentration was calculated using:

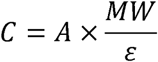

where A is absorbance at 280 nm, MW is the molecular weight of spike protein (540 kDa trimer equivalent), and ε is the extinction coefficient (428,225 ML¹ cmL¹).

Following concentration measurement, all enzyme-treated and mock-treated (as control) spike protein preparations were adjusted to a final concentration of 0.5 mg mLL¹ prior to downstream platelet activation experiments. Successful deglycosylation following PNGase F and sialidase treatment was verified by SDS-PAGE mobility shift and lectin blotting.

### Western and lectin blot analysis

Enzyme-treated and mock-treated Spike protein run through SDS-PAGE 10% Bis-Tris NuPAGE (Invitrogen, NW00105BOX), one set left for Coomassie blue staining, the rest of the set of gels transferred to nitrocellulose membrane (GE Healthcare). Membranes were blocked with 5% (w/v) skim milk in Tris Buffered Saline, 0.05% Tween20 (TBST) for western blot, washed and probed with anti-his monoclonal antibody. HRP conjugated anti-mouse (Dako, P0161) was used as secondary antibodies, and blots was imaged by chemiluminescence (SuperSignal TM, Thermo scientific, 34580). Western blots were imaged using BioRad ChemiDoc XRS+. Membranes were blocked with 3% BSA in TBST for lectin blot, and incubated with Sambucus nigra Lectin (SNA, FITC conjugated) for sialic acid detection, Wheat germ agglutinin (WGA, FITC conjugated) for the detection of sialic acid and N-acetylglucosamine, peanut agglutin (PNA, FITC conjugated) for the detection of galactose. Lectin blots were washed and imaged with using BioRad ChemiDoc XRS+ with 488 nm laser excitation.

### Statistical Analysis

Statistical analyses were performed using GraphPad Prism. Two-tailed Student’s t test (paired) and One-way ANOVA (paired) with Dunnett multiple comparison test were used in the data analysis. Statistical significance is expressed as p <0.05 (_∗_), p <0.01 (_∗∗_), p <0.001 (_∗∗∗_) for all tests.

## Supporting information

Supplementary Data for Bremaud et al

## Acknowledgments

General

We express our gratitude to Dr Oren Cooper at IBG, Griffith University, for generously providing us with lectins used in lectin blots in this study. EM analyses were performed at the Centre for Microscopy and Microanalysis at the University of Queensland (CMM UQ), Australia. We thank Drs Nicole Schieber, Kathryn Green, Erica Lovas, and Hui Diao at CMM UQ for their assistance and guidance in CLEM, FIB-SEM, and thin-section TEM set up.

## Author contributions

AB, BLS, and JM designed the study. AB, AS, EB, VM and BLS performed the experiments. LD and MvI contributed to reagents. AB and JM wrote the manuscript. All authors contributed to the discussions on the experiments, the collected data, and the finalisation of the manuscript.

## Data and materials availability

All data needed to evaluate the conclusions in the paper are present in the paper and / or in the Supplementary Materials. Additional data related to this may be requested from the corresponding author.

---

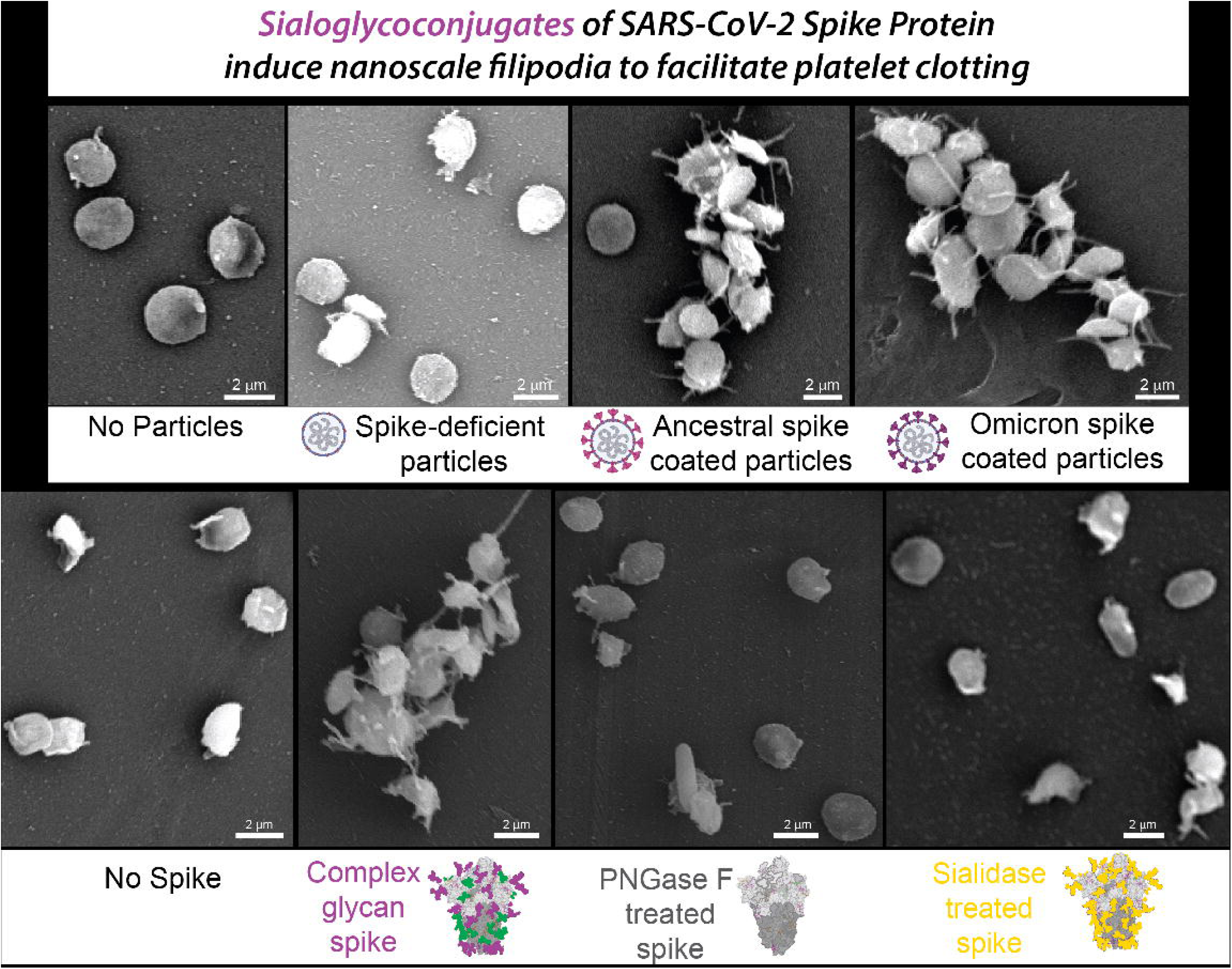

