## Supplementary Data for Bremaud et al for "SARS-CoV-2 spike protein-associated sialoglycoconjugates induce nanoscale filipodia to facilitate micro-size platelet clotting"

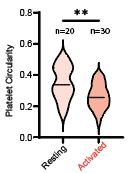

***Figure S1***. Quantitative comparison showed different circularity of resting- (light orange) and activated- (dark orange) platelets (Circularity values measured in ImageJ range from 0 to 1, where 1 represents a perfect circular (resting-like) platelet shape, and values closer to 0 indicate progressively irregular or spread platelet morphology associated with activation.).

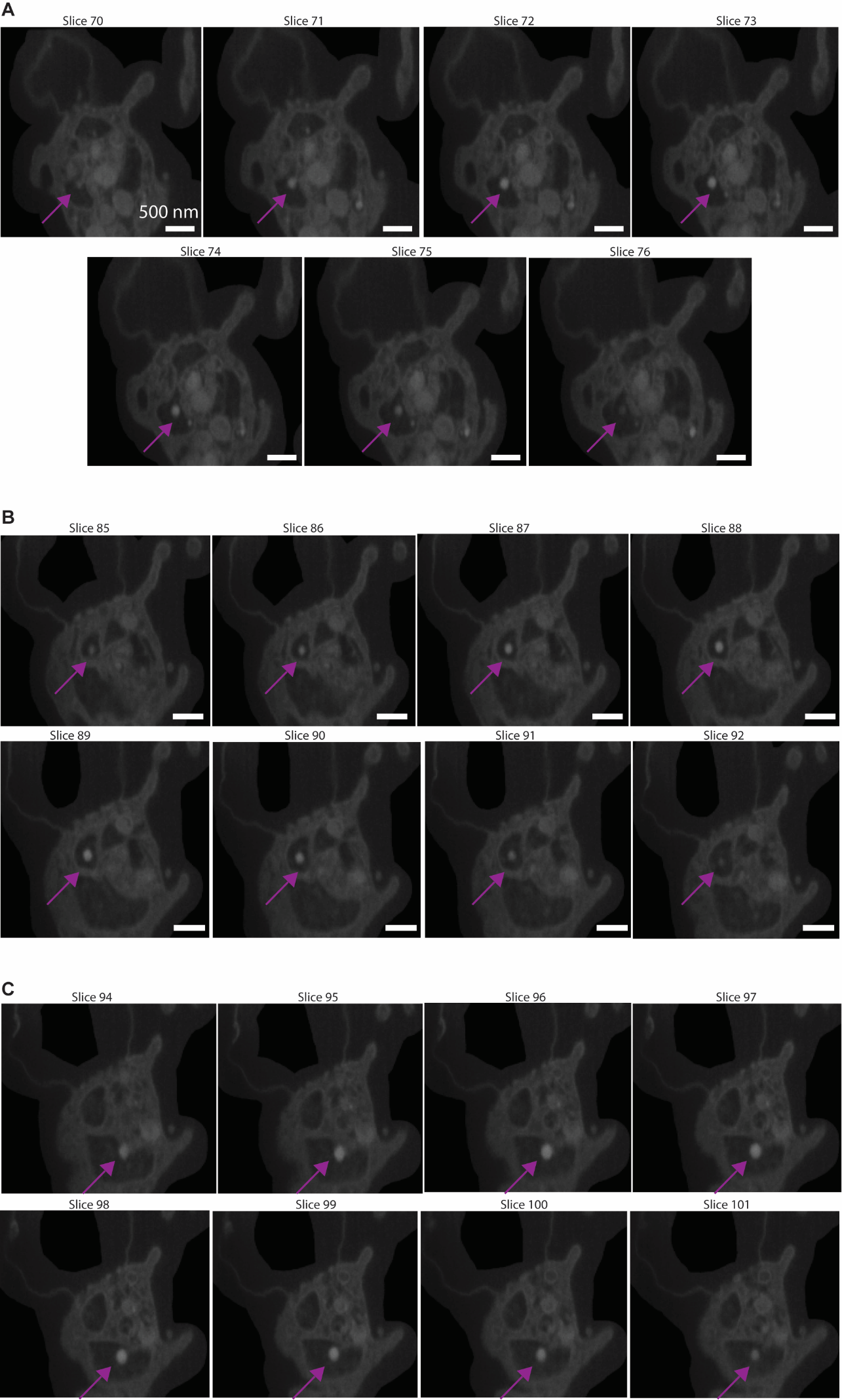

***Figure S2***. Slice-view examples showing omicron spike coated particles trapped within the open canalicular system (OCS) of platelets. (A–C) Three representative platelet examples demonstrating VLP localization within the OCS. For each example (A, B, and C), sequential slice-view images (7–8 slices , each slice thickness is 20 nm) are shown to illustrate the spatial continuity of the particles signal within the OCS channels across the z-direction (corresponding to particles sizes of approximately 140 nm), confirming internal association rather than surface attachment. Purple colour arrow indicates representative VLPs observed within the OCS structures.

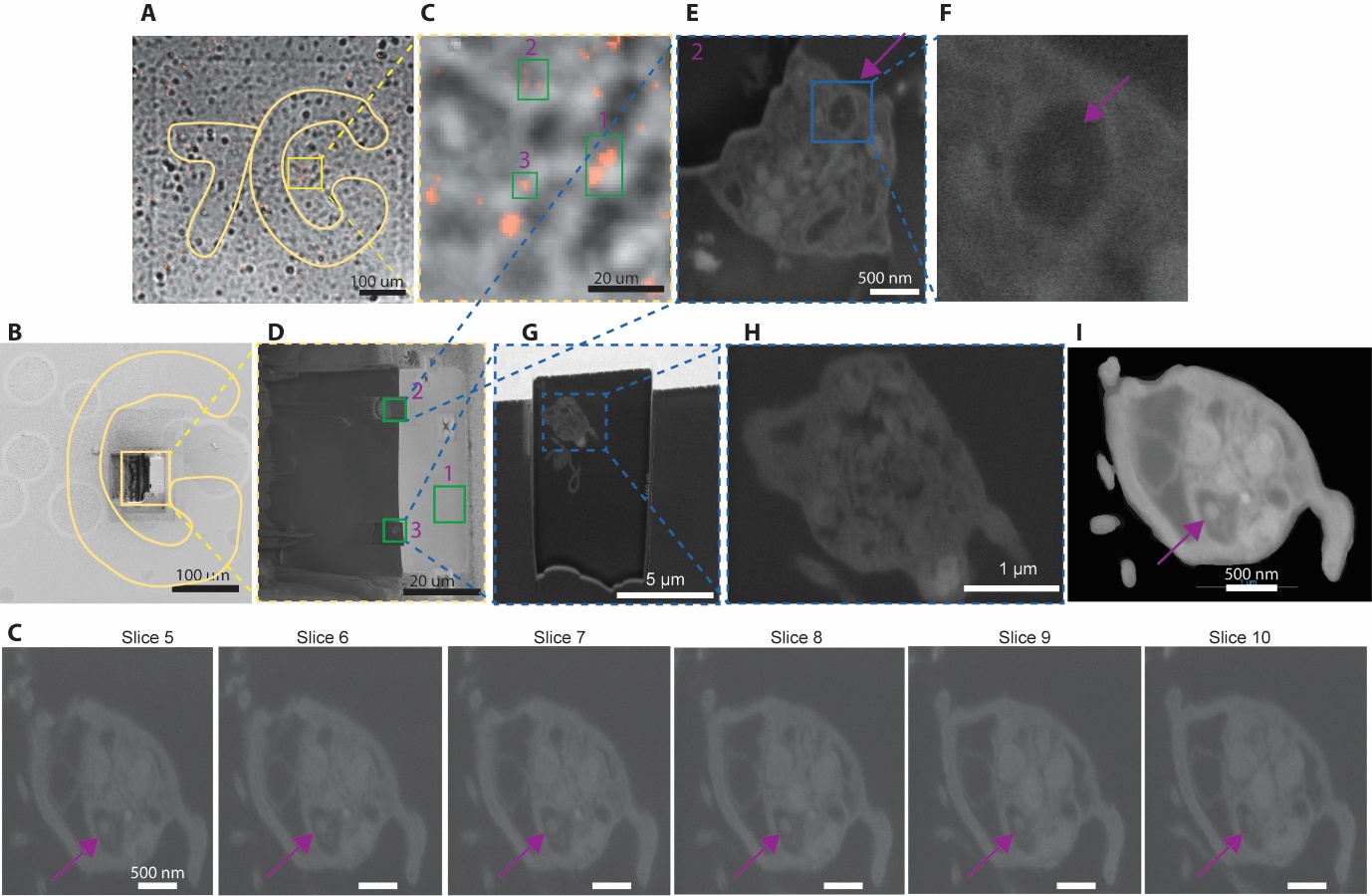

***Figure S3*.** Correlative confocal and FIB–SEM workflow for targeting regions of interest on a gridded coverslip. (A) Confocal microscopy image showing the alphanumeric grid reference (7G highlighted) on the coverslip used for region-of-interest localization. (B) Corresponding FIB–SEM image of the same grid region confirming successful coordinate correlation between confocal and electron microscopy. (C) Higher-magnification view of the selected region from (A), with three target areas (regions 1–3) marked for FIB milling. Region 1 is presented in the main manuscript. (D) FIB–SEM image of region 1 following milling. (E, F) FIB–SEM images of region 2 after milling. (G–H) FIB–SEM images of region 3 after milling, demonstrating successful targeting of multiple sites within the correlated grid location. (I) Contrast enhanced image (from raw images in J) of platelet showing particles trapped within OCS. (J) Representative platelet example demonstrating VLP localisation within the OCS. The sequential slice-view images from region 3 in D (6 slices, each slice thickness is 20 nm) are shown to illustrate the spatial continuity of the particles signal within the OCS channels across the z-direction (corresponding to particles sizes of approximately 120 nm), confirming internal association rather than surface attachment. Purple colour arrow indicates representative VLPs observed within the OCS structures.

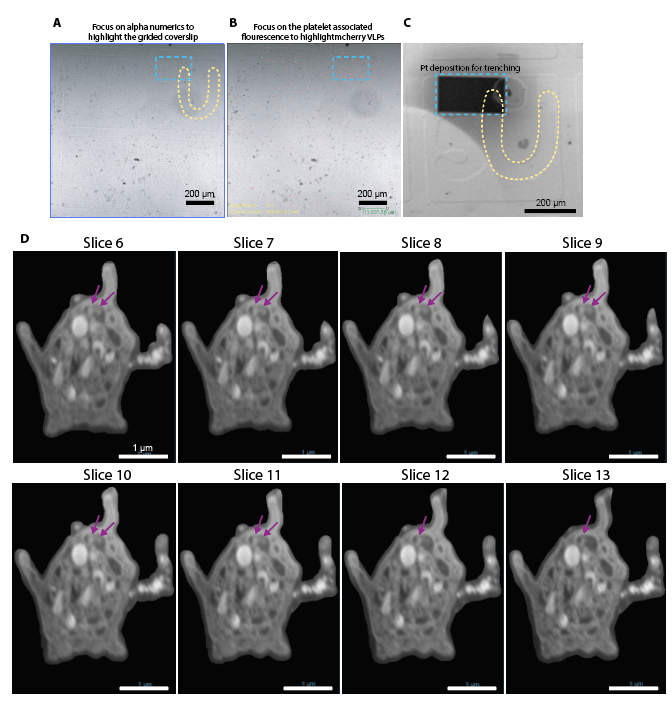

***Figure S4*.** Correlative confocal and FIB–SEM identification of mCherry-tagged omicron coated particles within the platelet open canalicular system (OCS). (A) Confocal microscopy image focused on the alphanumeric grid pattern on the coverslip used for coordinate registration. (B) Corresponding fluorescence image highlighting the platelet region of interest (ROI); the blue dashed square indicates the selected targeting area. (C) FIB–SEM image of the same ROI after relocation in the electron microscope, showing platinum (Pt) deposition over the target region prior to milling. (D) Representative slice-view sequence (8 consecutive slices) demonstrating two closely positioned VLPs within the platelet OCS, supporting their localization inside the canalicular network rather than on the platelet surface.

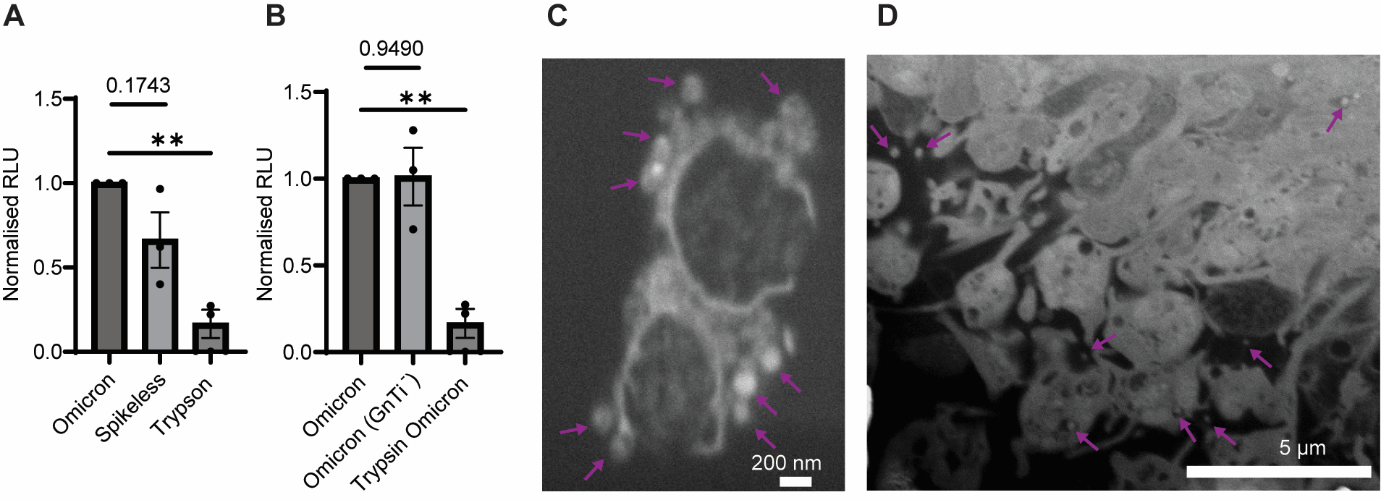

***Figure S5*.** Characterisation of spike protein-coated particles and their association with platelets. (A) Normalized relative luminescence units (RLU) measured from Omicron spike protein–coated particles, spike protein-deficient (spikeless) control particles, and trypsin-treated platelets and omicron spike protein–coated particles, confirming possible spike protein-dependent signal contribution and association of particles with platelets. (B) Normalized RLU comparison of Omicron spike protein–coated VLPs produced in standard cells and in GnTI⁻ cells to enrich high-mannose glycosylation on the spike protein surface, including corresponding trypsin-treated controls. (C, D) Representative FIB–SEM cross-sectional images showing spike protein-coated particles associated either within the platelet open canalicular system (OCS) or at the external platelet surface, illustrating multiple modes of particle–platelet interaction. (Purple arrows indicate particle positions relative to the platelet membrane and OCS structures).

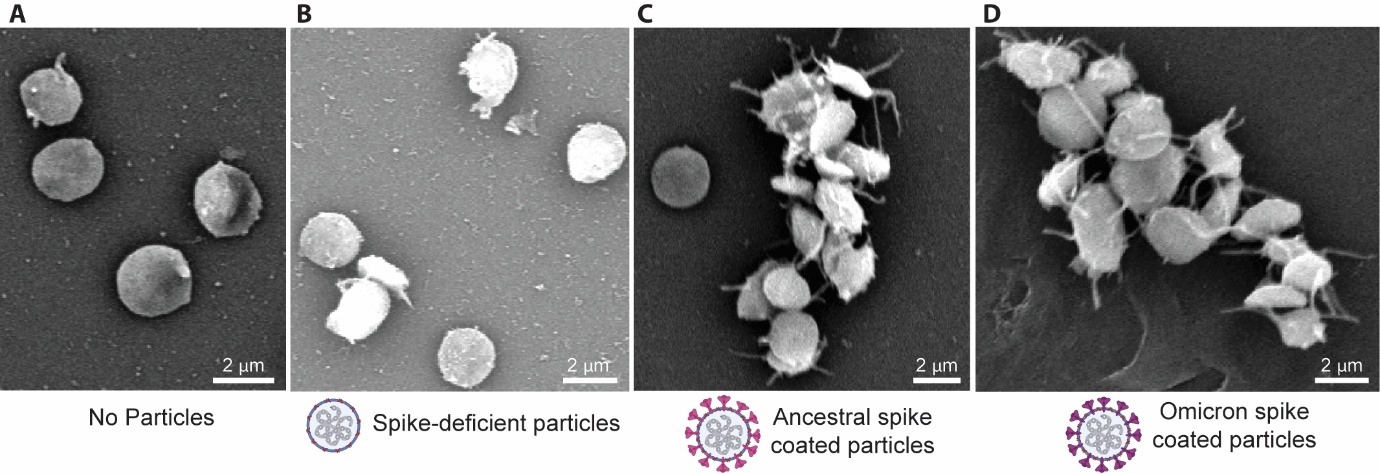

***Figure S6*.** Viral particle-dependent platelet activation visualised by scanning electron microscopy (SEM). (A) SEM image of platelets incubated without particles (negative control), showing predominantly dispersed platelets with minimal morphological features of activation, such as absent or very limited filopodia extrusion. (B) SEM image of platelets incubated with spike protein-deficient particles, displaying morphology comparable to the no-particle control in (A), with platelets remaining largely dispersed and exhibiting minimal filopodia formation. (C) SEM image of platelets incubated with ancestral spike protein-coated particles, demonstrating clear platelet aggregation and pronounced activation morphology, including extensive filopodia extrusion. (D) SEM image of platelets incubated with Omicron spike protein-coated particles, showing a phenotype like (C), with evident platelet aggregation and strong activation features characterised by prominent filopodia extension.
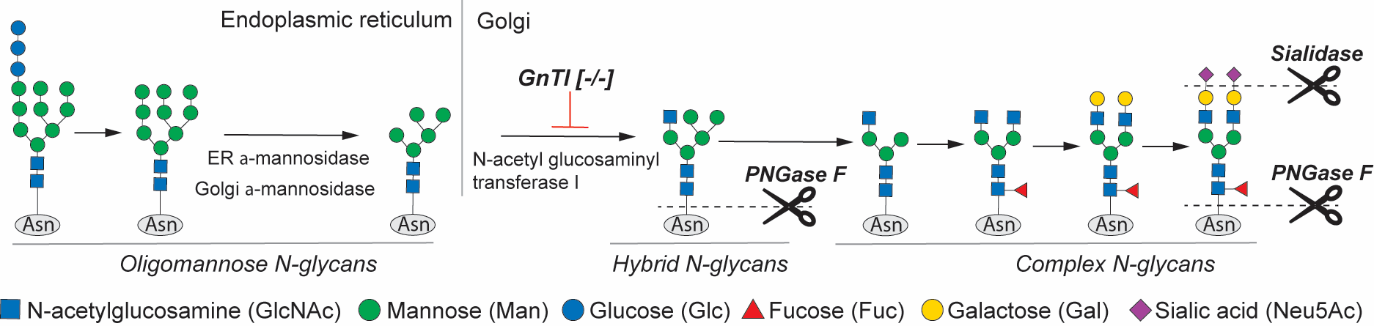

***Figure S7***. N-Glycan biosynthetic pathway. N-acetyl glucosaminyl transferase I (GNT1) action in the pathway, plus PNGase F and Sialidase enzyme cleavage sites are indicated on hybrid and complex glycans.

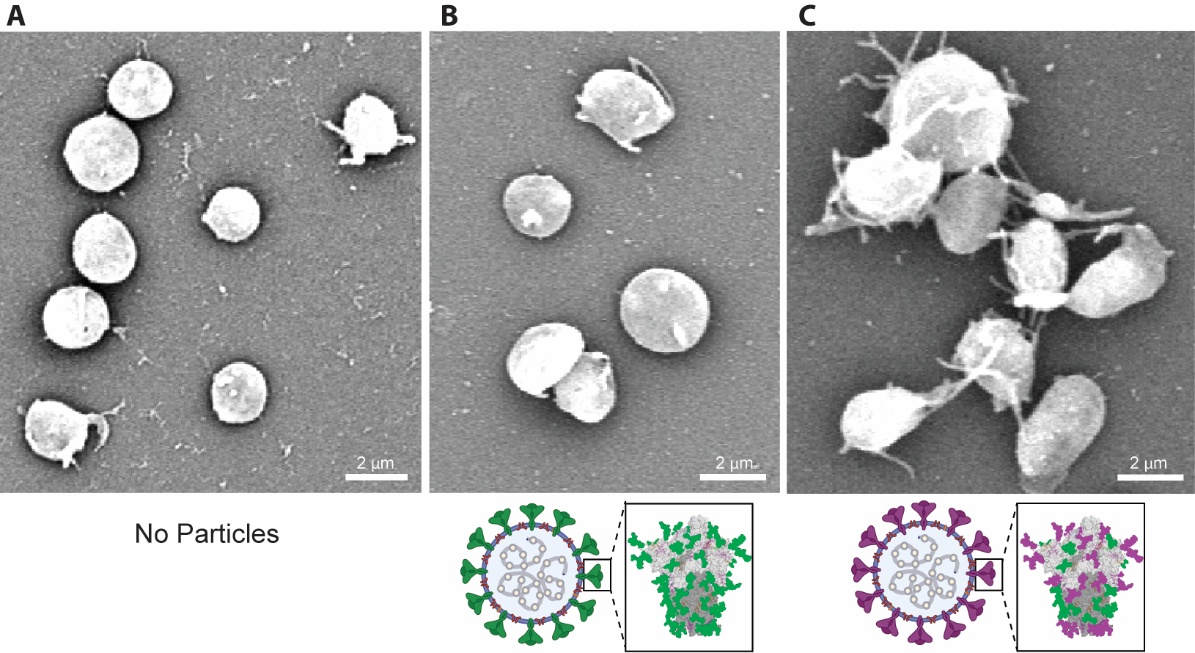

***Figure S8*.** Platelet activation depends on glycosylation pattern of Omicron spike protein–coated particles. (A) SEM image of platelets incubated without particles (negative control), showing predominantly dispersed platelets with resting morphology and no evident filopodia extrusion. (B) SEM image of platelets incubated with Omicron spike protein containing high-mannose glycoprotein–coated particles, demonstrating morphology comparable to the control in (A), with platelets remaining dispersed and lacking clear activation features such as filopodia extension. (C) SEM image of platelets incubated with Omicron spike protein containing sialic acid enriched complex glycoprotein–coated particles, showing pronounced platelet aggregation and clear activation morphology characterised by extensive filopodia extrusion.

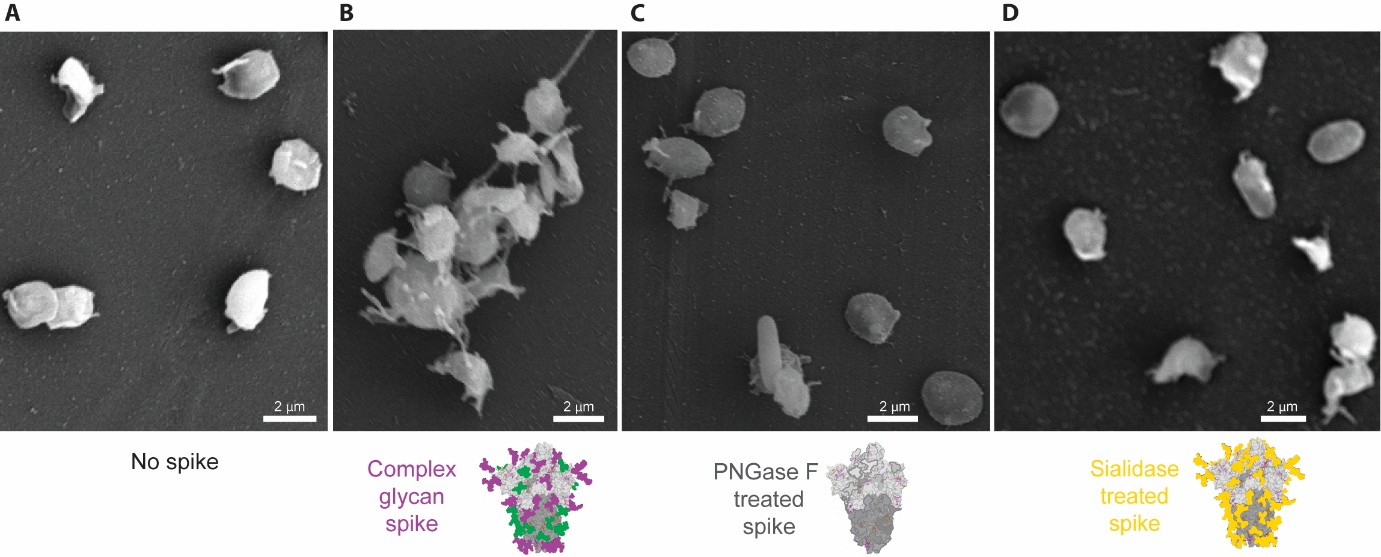

***Figure S9*.** N-linked glycosylation and terminal sialic acid residues on spike protein regulate platelet activation and aggregation. (A) SEM image of platelets incubated without spike protein (negative control), showing predominantly dispersed platelets with resting morphology and less evident filopodia extrusion. (B) SEM image of platelets incubated with terminal sialic enriched complex glycan–containing spike protein, demonstrating pronounced platelet aggregation and clear activation morphology characterised by extensive filopodia extrusion. (C) SEM image of platelets incubated with PNGase F–treated spike (N-linked glycans removed), showing markedly reduced platelet aggregation and minimal filopodia extrusion compared with (B), indicating reduced activation. (D) SEM image of platelets incubated with sialic acid free (or galactose terminal-capped) spike protein, displaying morphology like control conditions in (A) and PNGase F–treated samples in (C), with largely dispersed platelets and minimal filopodia formation.

***Supplementary Video S1*.** FIB–SEM slice-view imaging and three-dimensional volume reconstruction animation reveal the spatial organisation of particle in platelet OCS and interactions. Serial FIB–SEM sections were acquired and reconstructed into a 3D volume using Dragonfly software to visualise three aggregated platelets in close contact. Within a platelet, three Omicron spike protein–coated particles are clearly resolved and appear physically trapped inside the OCS.
